# Calibration of instrumented treadmills using an instrumented pole; a modified version to use relatively smaller forces

**DOI:** 10.1101/2024.07.08.602562

**Authors:** Seyed-Saleh Hosseini-Yazdi

## Abstract

The instrumented treadmills’ quality of the generated Ground Reaction Forces (GRF) may degrade over time, as the original calibration matrix may not accurately represent the exerted forces. A cost-effective alternative to manufacturer recalibration is to use an instrumented pole for calibration. Collins et al. presented a simple method to collect multiple data points by exerting forces in various directions. The sensor on the instrumented pole provides instantaneous force magnitudes, while motion capture records the pole’s instantaneous directions. They recommended a relatively large force magnitude (1000N), requiring at least two individuals. Using an optimization method, the new calibration may be estimate by relating the exerted forces (pole) to the treadmill signals. Here, we attempted to simplify the process further, allowing a single individual to perform force exertion with additional force exertion direction. Thus, the calibrating forces were reduced to one-third of the prior recommendation. This maintained the structural integrity of the pole and helped avoid inducing bending moments that could affect calibration results. The cross-validation score for test data (prediction score) was at least 0.92. Additionally, comparing GRFs in posterior/anterior and vertical directions with a benchmark treadmill for even walking revealed an average cross-correlation of 0.97.

## Introduction

The instrumented treadmill is widely used for recording ground reaction forces (GRF) in gait analysis applications, often combined with motion capture data for inverse dynamics and other analyses ^1^. The treadmill generates electrical signals representing the captured forces and moments through load cells. These electrical signals are converted into forces and moments using a transformation matrix. The assumption is that the signals are independent, so the transformation matrices provided by equipment manufacturers are usually diagonal.

Over time, the original calibration of treadmills may need to be revised, necessitating attention to the load cells. The common way to calibrate an instrumented treadmill is to use known dead weights to compare the output signals ^2–4^. However, this method only allows for vertical load calibration. Collins et al. ^5^ proposed a simplified method that uses an instrumented pole to exert forces on the instrumented treadmill’s surface. The magnitude and direction of these forces were determined by recording the positions of markers affixed to the pole (motion capture) and the load cell signals (Figure 1A).

**Figure 1:**
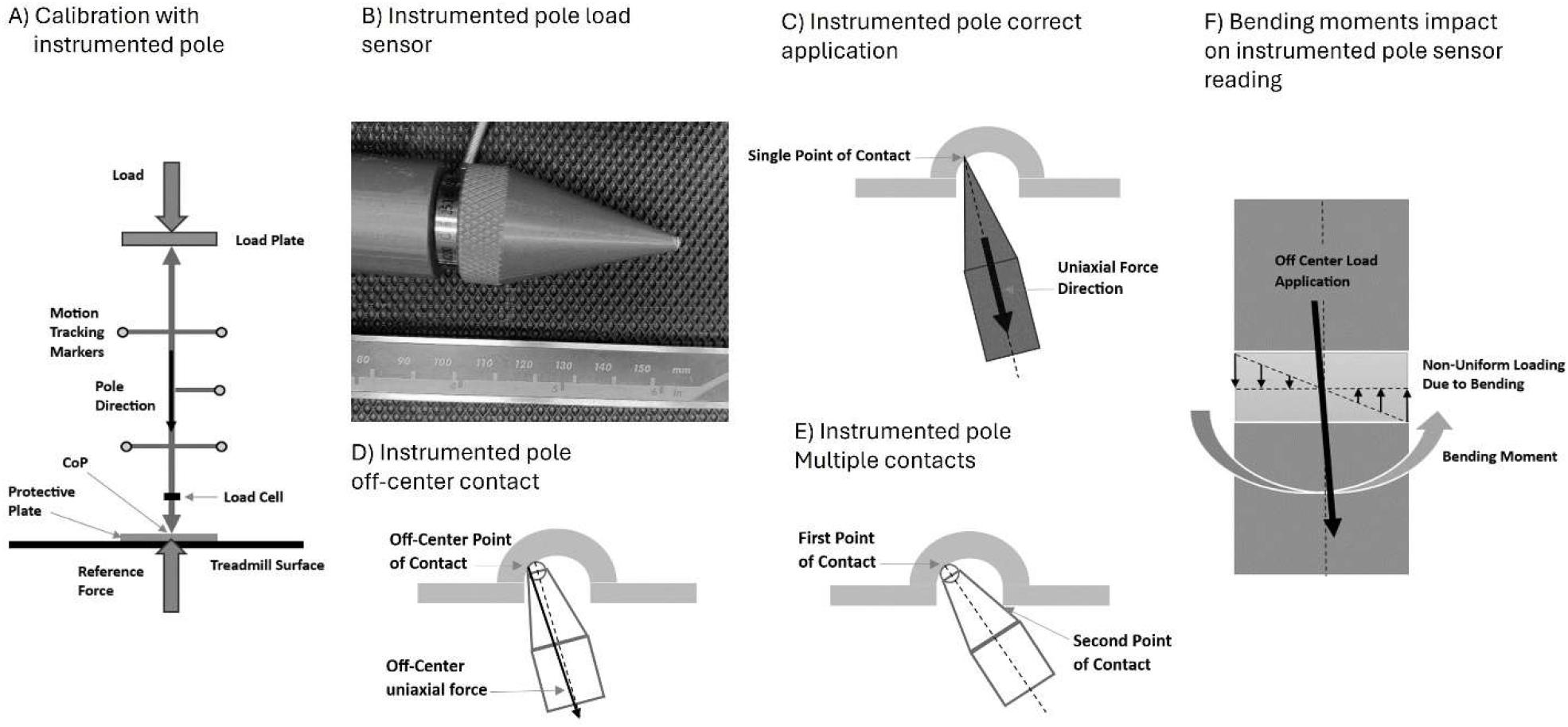
(A) Schematics depicting the load placement in arbitrary directions using an instrumented pole. Motion capture records the position and orientation of the pole during force exertion. (B) Close-up view of the load sensor integrated into the instrumented pole. (C) Illustration demonstrating the recommended method for generating uniaxial force using the instrumented pole. (D) Example of generating an off-center load, particularly relevant when the tip of the pole is worn out. (E) Depiction of multiple points of contact between the pole and handle during the force exertion process. (F) Potential consequence of deviating from the preferred force exertion method, leading to the generation of bending moments in the load gauge of the instrumented pole.

The force exertion in any direction ^6^ can be used to determine the reference Center of Pressure (COP) and the exerted forces ^7^. Collins et al. ^5^ calculated the associated moments about three axes by collecting force exertion data at 20 points along each treadmill belt. However, the authors observed differences in load signals when the same load was applied at different locations. Therefore, to ensure accurate calibration, they used two individuals to exert forces comparable to normal walking, as lower forces resulted in higher errors. They used the thousands of data points collected to map the treadmill’s signals to actual forces through an optimization method.

The instrumented poles used for treadmill calibration have conical tips to ensure that purely axial forces are generated by the load plate pushing to be captured by the pole’s load sensor (Figure 1 B&C). Applying loads other than through the tips can introduce bending moments (Figure 1D to F), leading to noisy readings. For instance, when two individuals attempt to exert large forces, the top tip may get trapped in the load plate seat. Additionally, excessive force application can wear out the pole tips, compromising the axial load’s purity. Such load exertions and estimation inaccuracies can reduce the accuracy of the derived transformation matrix.

In the following sections, we present a simplified version based on the original method ^5^. This method facilitates the calibration of instrumented treadmills and force plates and allows for force exertion by a single individual.

## Materials and Methods

To conduct uneven walking experiments, we modified an instrumented treadmill (Bertec, Columbus, OH, USA) by vertically extending the belt support frame structure to create clearance between the treadmill belt and the underlying support structure. This process involved releasing the top structure assembly by opening the load cell collars (Figure 2A). After component replacement, we reinstalled the top structure assembly and secured the load cell collars following the manufacturer’s procedure.

**Figure 2:**
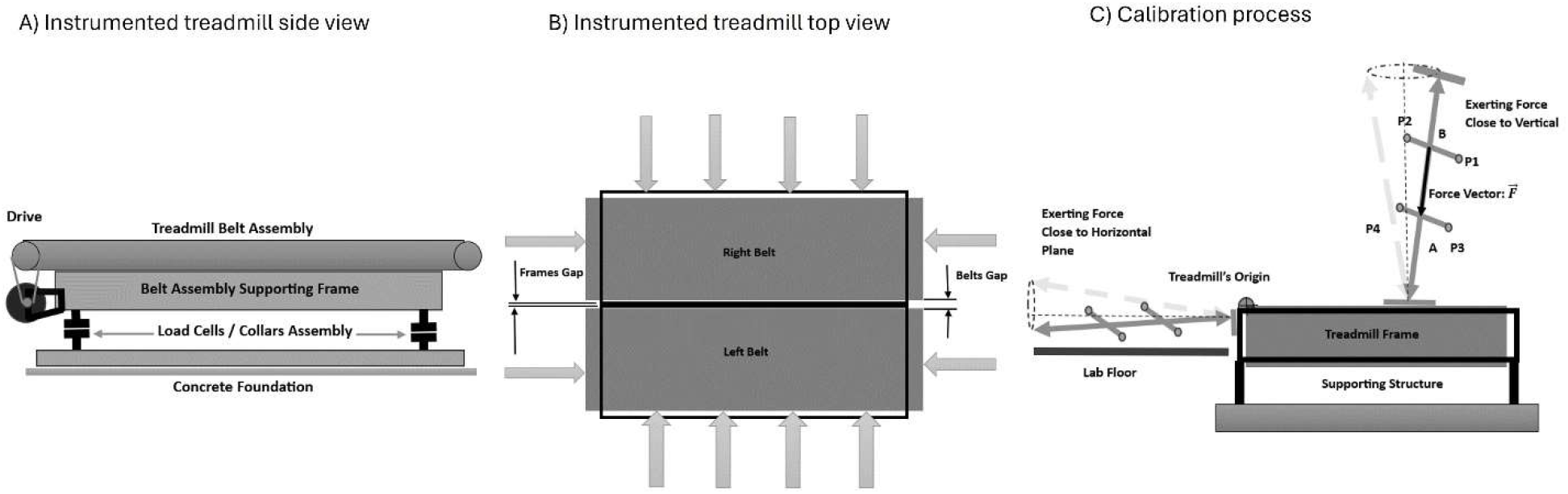
(A) Side view of an instrumented treadmill exhibiting the load cells, which generate force signals for any load placement. (B) Illustration depicting additional load exertions near the horizontal plane to compensate for lower calibrating forces. (C) demonstrating the force exertions close to both horizontal and vertical planes.

Preliminary trials indicated that an average person could exert a maximum uniaxial force of around 450 N. On the other hand, during a typical walking trial, an average person with a body mass of 75 kg may create vertical peak forces exceeding 800 N. Similarly, the Posterior/Anterior (P/A) force peaks may exceed 200 N. Therefore, one person’s force exertion would be nearly half the recommended magnitude ^5^. We initially decided to exert forces along the instrumented treadmill’s known axis to reduce noise associated with lower calibrating forces. However, to avoid generating zero forces or moments in one or more directions, we applied forces in directions that made small angles with the treadmill’s axes ^5,6^. When forces were used close to the horizontal plane, we disregarded the weights of the pole and plates, while in vertical positions, we added those weights in our calculations. We followed the original method for the number of force exertion instances on top of the treadmill belt ^5^ and added more instances to collect data close to the horizontal plane (Figure 2B).

To record the direction of the instrumented pole ^8^, we affixed passive markers to four pole handles (sampling rate 240 Hz, Optitrack Motion Capture system; NaturalPoint Inc. Corvallis, OR USA). We derived the positions of two points (upper: B and lower: A) as averages of marker positions to calculate directional cosines representing the force exertion direction (Figure 2C).

As the instrumented pole’s signal was recorded at a 960Hz, motion capture data was up-sampled. Each recording lasted for 10 seconds. Whenever the instrumented pole’s data recording was triggered (ON or OFF), a signature signal was sent to the motion capture system as a one-shot rising signal. Only the motion capture data recorded between the signature signals was used for analysis.

We divided the recorded data into training and test groups. The training data included both vertical and horizontal force exertion samples, while for testing, we used only vertical force exertion samples. We derived the transformation matrices for the right and left belts using the linearized model ^5^ based on the training data set. Accordingly, we derived scores for actual and predicted force and moment values to quantify prediction quality (sci-kit-learn 1.3.2).

## Results

The resulting transformation matrices exhibited similarities to the manufacturer’s proposal, with large values in diagonal elements. However, unlike the manufacturer’s, these matrices contained off-diagonal elements^5^. Based on the training data, the average prediction scores for the left and right belts were 0.9844 ± 0.0134 and 0.9925 ± 0.0035, respectively. A visual inspection of estimated forces and moments against reference values for the testing data also indicated a close tracking. The average prediction scores for the testing data were 0.9228 ± 0.0708 and 0.9333 ± 0.0759 for the left and right belts, respectively (Figure 3A). We compared the GRF of even walking at 1.2 m · s^-1^ (duration: 60 s) on the calibrated treadmill and compared it with a similar machine (benchmark). The cross correlation and Bland and Altman agreements ^9^ scores were 0.97 and 96% respectively. The subject’s mass was also estimated by averaging the total vertical GRF throughout data collection. The subject’s mass estimation by the calibrated treadmill was 0.46% higher than the benchmark’s (73.04 kg versus 73.37 kg, Figure 3B).

**Figure 3:**
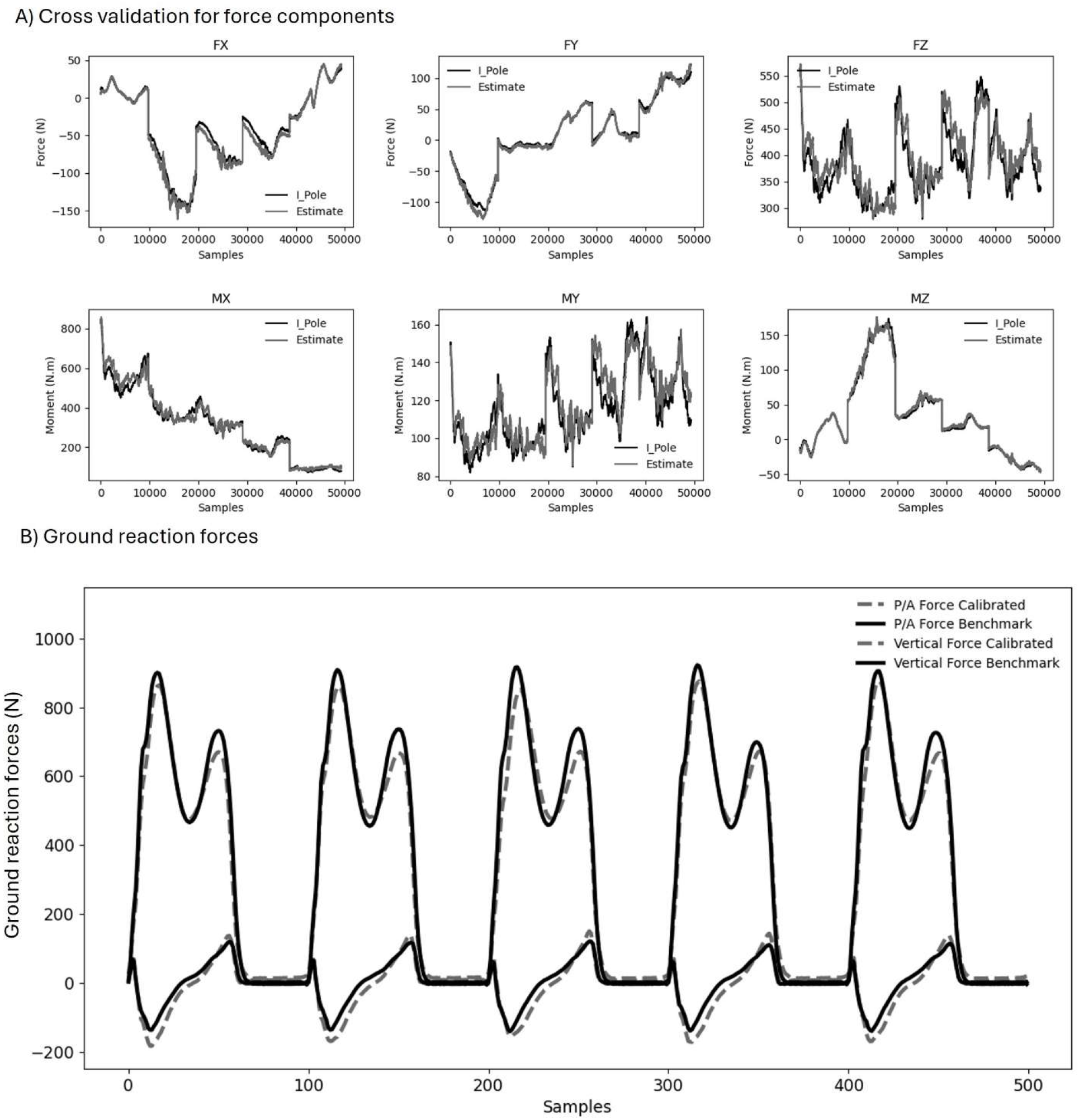
(A) Cross-validation results for the six components of the test data show prediction scores exceeding 0.92 for both the right and left belts. (B) A comparison of the ground reaction forces (GRF) during even walking between the benchmark and calibrated treadmill reveals a correlation score of 0.97.

## Discussions

We calibrated an instrumented treadmill structurally modified to accommodate uneven walking experiments terrains. This modification presented additional weight, which may have affected the load cells’ original readings. Consequently, we adjusted the load cells’ shimming to ensure consistent force signal generation for the same weights. We successfully calibrated the instrumented treadmill with modifications to enable a single-individual force exertion for the process. This approach not only preserved the structural integrity of the instrumented pole, but also eliminated calibration inaccuracies associated with lower forces applications.

The resulting transformation matrices exhibited some differences from the manufacturer’s matrices. Extended usage, structural compromises, and weight additions from the modification likely contributed to these deviations. The prediction score for the testing data was slightly lower than that for the training data (average of 0.0604 declines), with increased variations (averages: 0.0733 versus 0.0085). Nevertheless, the high prediction score (0.9281) for the testing data suggests we can confidently use the derived transformation matrices for our experiments. The slight difference in prediction scores between testing and training data cannot be attributed to data leakage, as the data for testing and training were kept separate. We also ensured that the pole’s Center of Pressure (COP) did not coincide with the locations of the training data. The test data size is comparable to typical model training and evaluation data ratios (test-to-train sample data ratio: 1:5). Both testing and training data were collected on the same day with the same instruments and underwent the same preprocessing. Therefore, we cannot also attribute additional noise induction in the testing data. While the test data only consisted of near-vertical collections, it also included horizontal force components, suggesting that the training and test data distributions were similar. The close prediction scores indicate that overfitting to the training data likely did not occur.

Based on the results obtained from the Bland-Altman analysis and cross-correlation tests, it can be inferred that there exists a high degree of similarity between our calibrated and the benchmark treadmills as intended for the walking trials. These scores suggest a strong agreement between the two readings, indicating a consistent relationship and alignment in their measurements. Such findings reinforce the notion that both treadmills exhibit comparable performance characteristics.

In summary, extensive use or any modifications to instrumented treadmills can potentially compromise the original calibration. However, the use of an instrumented pole allows for force exertion in various directions and magnitudes ^5^, making it a cost-effective and readily available tool. This approach provides an abundance of data points to train the linear model and derive accurate calibrations for the instrumented treadmill. While it has been suggested that manual force exertions may not yield reproducible calibration results ^10^, the high volume of test data and strong testing and correlation scores indicate the reliability of the modified method used.

## Acknowledgements

This work was supported in part by the Natural Sciences and Engineering Research Council of Canada (NSERC) Discovery and Canada Research Chair (Tier 1) programs.

